# Effects of non-invasive brain stimulation on visual perspective taking: A meta-analytic study

**DOI:** 10.1101/2021.04.24.441219

**Authors:** Yuan-Wei Yao, Vivien Chopurian, Lei Zhang, Claus Lamm, Hauke R. Heekeren

## Abstract

Visual perspective taking (VPT) is a critical ability required by complex social interaction. Non-invasive brain stimulation (NIBS) has been increasingly used to examine the causal relationship between brain activity and VPT, yet with heterogeneous results. In the current study, we conducted two meta-analyses to examine the effects of NIBS of the right temporoparietal junction (rTPJ) or dorsomedial prefrontal cortex (dmPFC) on VPT, respectively. We performed a comprehensive literature search to identify qualified studies, and computed the standardized effect size (ES) for each combination of VPT level (Level-1: visibility judgment; Level-2: mental rotation) and perspective (self and other). Thirteen studies (rTPJ: 12 studies, 23 ESs; dmPFC: 4 studies, 14 ESs) were included in the meta-analyses. Random-effects models were used to generate the overall effects. Subgroup analyses for distinct VPT conditions were also performed. We found that rTPJ stimulation significantly improved participants’ visibility judgment from the allocentric perspective, whereas its effects on other VPT conditions are negligible. Stimulation of dmPFC appeared to influence Level-1 performance from the egocentric perspective, although it was only based on a small number of studies. Notably, contrary to some theoretical models, we did not find strong evidence that these regions are involved in Level-2 VPT with a higher requirement of mental rotation. These findings not only advanced our understanding of the causal roles of the rTPJ and dmPFC in VPT, but also revealed the efficacy of NIBS on VPT is relatively small. Researchers should also be cautious about the potential publication bias and selective reporting.

## 1 Introduction

The ability to take another’s perspective is crucial for navigating complex social environments. To view the world from the second-person standpoint requires that one distinguishes between the self and the other in relation to the environment (Kessler and Rutherford, 2010; Lieberman, 2007). One social cognitive process that is closely related to this ability is visual perspective-taking (VPT). Dysfunction related to VPT has been observed in multiple clinical disorders, including autism and schizophrenia (Eack et al., 2017). Thus, it is essential to identify cognitive and neural mechanisms underlying the VPT process, as a steppingstone to target interventions for related disorders.

Flavell and colleagues (1977; 1981), identified two levels of VPT. Level-1 VPT refers to the ability to judge an object’s visibility from the perspective of both the self and other. Consider, for example, playing hide-and-seek: You need knowledge about what the other person can see to hide from them. Children around the age of 18-24 months (Flavell et al., 1981) as well as chimpanzees (Braeuer et al., 2007), dogs (Hare and Tomasello, 2005) and goats (Kaminski et al., 2005) show the ability to make such line-of-sight judgements. Level-2 VPT, on the other hand, enables humans to describe how an object looks from another’s perspective and establishes a shared view of the world by creating a common reference frame for spatial localizations (Flavell, 1977; Kessler and Rutherford, 2010; Michelon and Zacks, 2006). For instance, imagine standing in front of a car, while your friend views it from behind: you are aware that although the car is visible to both of you, your friend has a different visual perspective on it (Pearson et al., 2013). Thus, Level-2 VPT has a higher level requirement of embodied rotation compared to its Level-1 counterpart (Martin et al., 2020).

In recent years, researchers have conducted a few neuroimaging studies to assess the neural mechanisms underlying VPT. One candidate region identified for this process is the temporoparietal junction (TPJ), as both the right and left parts of this region appear to play a critical role in multiple processes relevant to VPT, including detecting self-other incongruences, controlling self and other representations, and inhibiting the influence of the non-relevant representation via orienting attention (Bahnemann et al., 2009; Lamm et al., 2016; Quesque and Brass, 2019; Wolf et al., 2010). Indeed, the bilateral TPJ has often been reported across different VPT conditions (Bukowski, 2018; Schurz et al., 2013). Another critical region for integrating self-other processing is the dorsomedial prefrontal cortex (dmPFC). The dmPFC has also been implicated in making judgements about others (Denny et al., 2012), social information processing (Lieberman et al., 2019), and in introspection and assessment of mental states (Dore et al., 2015). In VPT tasks, the dmPFC has been reported when requiring egocentric perspective taking and suppressing the influence of the other’s perspective, with a proposed process of imagining movement and suppressing the motor response to physically rotate the body (Bukowski, 2018; Mazzarella et al., 2013; Munzert et al., 2009).

While these neuroimaging studies highlight candidate brain hubs for self-other differentiation and integration in VPT, they are mostly based on correlational methods and thus causal relationships remain to be established (Bell and DeWall, 2018; Lieberman et al., 2019). Fortunately, non-invasive brain stimulation (NIBS) techniques, including transcranial direct current brain stimulation (tDCS) and transcranial magnetic stimulation (TMS), provide an approach to overcome this limitation (Donaldson et al., 2015; Polania et al., 2018). Specifically, tDCS applies weak direct currents to cortical regions. It could either facilitate or inhibit the spontaneous neuronal activity depending on the polarity of the electrode. Typically, anodal and cathodal stimulation has been shown to increase and decrease cortical excitability, respectively (Bell and DeWall, 2018; Brunoni and Vanderhasselt, 2014). TMS, on the other hand, uses a changing magnetic field to induce an ionic current at a brain region based on the principle of electromagnetic induction. The effects of TMS depend on factors including frequency, intensity, and duration of stimulation. For example, single-pulse TMS could depolarize the targeted neurons, whereas high-frequency (e.g., >10 Hz) repetitive TMS (rTMS) typically disrupts the cortical function during the stimulation (Kobayashi and Pascual-Leone, 2003).

Researchers have increasingly used NIBS to investigate the causal role of different brain regions in VPT in the past few years. For example, anodal tDCS of the right TPJ (rTPJ) has been shown to improve participants’ performance when judging an item’s visibility from another’s perspective (Santiesteban et al., 2012). Moreover, another study showed that such an improvement could extend to Level-2 allocentric perspective-taking (Martin et al., 2019a). However, there is also opposing evidence in which the rTPJ stimulation increased the impact of perspective discrepancy during Level-1 VPT (Martin et al., 2020). Thus, despite that much effort has been devoted to clarifying the causal relationships between brain regions and VPT, the overall findings to date paint a rather mixed and inconclusive picture.

The inconsistency may be partly due to the complexity and heterogeneity of the existing VPT paradigms (Bukowski, 2018). As mentioned above, participants may be asked to judge the visibility or location of a target from different perspectives (e.g., self or other), in which distinct underlying cognitive mechanisms may be involved. For example, Level-2 VPT typically requires more embodied processing than Level-1 VPT (Martin et al., 2020). However, there is no consensus yet if stimulating a brain region would selectively influence any VPT conditions. Moreover, with a few exceptions (Martin et al., 2019b,a, 2020), most NIBS studies in this field only focused on one brain region, making it difficult to compare the different regions’ roles in VPT.

The current study aims to clarify the causal roles of key brain regions in VPT. Based on the feasibility of the included studies, we focused on studies targeting rTPJ and dmPFC and quantitatively synthesized the effects of stimulation of these two regions on distinct VPT components.

## 2 Methods

The meta-analysis was conducted following the Cochrane Handbook for Systematic Reviews of Interventions (Higgins et al., 2019) and PRISMA guidelines for meta-analyses (Liberati et al., 2009). The literature search and review, as well as data extraction, were performed by two co-authors (Y.W.Y and V.C) independently. Discrepancies were resolved by discussion.

### 2.1 Search strategy and eligibility criteria

An online literature search was conducted in Pubmed, Web of Science, and ProQuest for full-text articles from January 2000 to June 2020 without language restrictions. The following query syntax was used: (“stimulation” OR “TMS” OR “tDCS” OR “tACS” OR “tPCS” OR “tRNS” OR “TBS”) AND (“perspective taking” OR “perspective-taking” OR “VPT”). To be included in the final meta-analysis, studies had to: (1) perform NIBS, (2) include a VPT task, (3) enroll healthy participants, (4) have a control or sham condition. Studies without full-text available were excluded. Note that, although previous brain stimulation studies mainly focused on rTPJ or dmPFC, we did not explicitly include “rTPJ” OR “dmPFC” during the literature search. However, as a random-effects model requires at least 3 effect sizes (ESs), the sample size limitation did not allow us to perform a meta-analysis on studies that targeted other brain regions.

We first identified 27 potentially related studies by checking the title and abstract. Two authors then independently decided if these studies should be included in the review by reading the full text. The inter-rater reliability for the article selection shows a high agreement (Cohen’s Kappa = 0.87, *p* < 0.001). The authors resolved their disagreement about two articles by discussion.The details were listed in Table S1. Finally, a total of 13 studies targeting rTPJ or dmPFC were included in the meta-analyses.

### 2.2 Quality assessment

We used the Cochrane risk of bias tool to assess the quality of studies. Ratings (low, high, or unclear risk of bias) were assigned to each study based on the following six criteria: (1) assessments for sequence generation, (2) allocation concealment, (3) blinding of participants and researchers, (4) blinding of outcome assessment, (5) incomplete outcome data, and (6) selective reporting.

### 2.3 Data extraction

For each included study, we extracted information regarding the sample size, age, and sex ratio. For intervention characteristics, we extracted the type of NIBS technique, stimulation region, blinding protocol, intensity, duration of active stimulation, valence (excitatory or inhibitory), and study design. For tasks, we extracted the VPT Level (1 or 2) and Perspective (Self or Other) information for each effect and focused on these four conditions in the following analyses.

As most VPT tasks had an experimental condition, where the object being judged was incongruent from the Self compared to the Other perspective, and a control condition, where the object was congruent from both perspectives, we extracted the means and standard deviation (SDs) of the congruency effect (i.e., incongruent-congruent for RT or congruent-incongruent for accuracy) for each VPT condition whenever possible. If there was no VPT control condition, we extracted the means and SDs of RT or accuracy of the incongruent trials. If the data were only presented in figures, means and SDs were estimated using the WebPlotDigitizer (Rohatgi, 2020). If only the standard error (SE) was available, we calculated the SD with the formula: 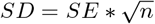

We also contacted the authors for the data and related information that was not reported, such as the correlation between repeated measures.

### 2.4 Data analysis

Data analysis was performed using R (version 4.0.1) and the Metafor package (Viechtbauer, 2010). We used the means and SDs to calculate the standardized ESs for each of the four conditions (VPT Level: 1 or 2, Perspective: Self or Other) and each stimulation target (rTPJ, dmPFC), respectively. For between-subject studies, we calculated the standardized mean difference (i.e., Hedge’s g). For within-subject studies, we calculated the standardized mean change (Morris and DeShon, 2002). We contacted the authors to ask for the raw data or correlation between repeated measures if the information was not provided in studies.

If a study reported both RT and accuracy, we calculated a combined ES and variance using the following equations:

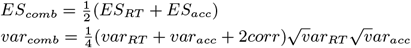

the correlation between RT and accuracy was not provided in studies, we used an assumed corr = 1, which was a conservative approach according to Borenstein et al. (2009) and Scammacca et al. 2014.

For three studies (Martin et al., 2020; Wang et al., 2016) that reported effects for different rotation angles and body postures during VPT, we focused on the 160-degree condition and calculated a combined ES for both body postures. One study (van Elk et al., 2017) that examined the effects of complex mental body transformation and stimulation sessions (online and offline) on VPT, to ensure its comparability with other studies, we only focused on the z-axis 180-degree condition and combined the ESs of both stimulation sessions. Moreover, the Australian group of Martin et al. (2019b) was also used in Martin et al. (2019a), so we only used the data from the South-East Asian group for Martin et al. (2019b).

For consistency, the direction of the ES was defined as positive if the excitatory stimulation increased the incongruent RT or decreased the incongruent accuracy, and negative for the inhibitory stimulation. For between-subject tDCS studies that included anodal, cathodal, and sham stimulation groups, only the anodal > sham comparison was used for the main meta-analysis to avoid the data of the sham group being used repeatedly. It should be noted that the anodal stimulation group of Martin et al. (2019a) is a subset of Martin et al., 2019a, so we only included sham > cathodal comparison for this study.

We first performed two separate meta-analyses to examine the overall effect of rTPJ and dmPFC stimulation on VPT, respectively. As some studies included multiple VPT conditions, we used both the two-level (first level: ES, second level: VPT condition) and three-level (third level: study) random-effects model with restricted maximum-likelihood estimator (Cheung, 2014; Konstantopoulos, 2011). A critical difference between the two models is that the former ignored the within-sample variance and treated ESs from different VPT conditions as independent, whereas the latter accounted for potential dependence between ESs from the same study. Model comparison based on Akaike’s information criteria (AIC) was conducted to test which model was better given the data. Heterogeneity among the included ESs was assessed using the Q and I2 tests. The funnel plot and Egger’s test was used to assess publication bias (Egger et al., 1997). If an Egger’s test revealed significant publication bias, the trim-and-fill method (Duval and Tweedie, 2000) was used to generate a corrected estimate after accounting for the effects of unpublished studies.

To investigate if rTPJ and dmPFC stimulation selectively influence any VPT conditions, we further conducted subgroup analyses for each of the conditions and each stimulation target (except for rTPJ stimulation on VPT Level-2 Self condition because of insufficient ESs). Q test was used to statistically compare the aggregate ESs of different subgroups.

To assess the reliability of the results, we conducted a few control analyses. First, we used a leave-one-study-out analysis to examine the influence of individual ESs. Second, we conducted a control analysis for the dependent variable measure by replacing each of the combined RT and accuracy ESs with ESs for only RT or accuracy and comparing the effect sizes. In addition, we explored the effects of stimulation timepoint (offline or online), study design (between- or within-subject design), and tDCS electrode size on results for TPJ stimulation. For studies using dmPFC stimulation, we examined the effects of contrast selection (cathodal or anodal) on results, because the studies were otherwise similar in their design.

## 3 Results

A total of 15 studies met the inclusion criteria for the qualitative review (Fig 1). Since a random-effects model requires at least 3 effect sizes (ESs), 3 studies were excluded from the meta-analysis because they stimulated regions not tested for in other studies and thus did not provide sufficient ESs required by a meta-analysis. Among the remaining 13 studies, 9 stimulated rTPJ only, 1 stimulated dmPFC only, and 3 stimulated both regions (see Table 1 for study overview). After distinguishing 4 unique combinations of VPT Level and Perspective, we obtained 23 ESs for the rTPJ and 14 ESs for the dmPFC targets.

**Figure 1:**
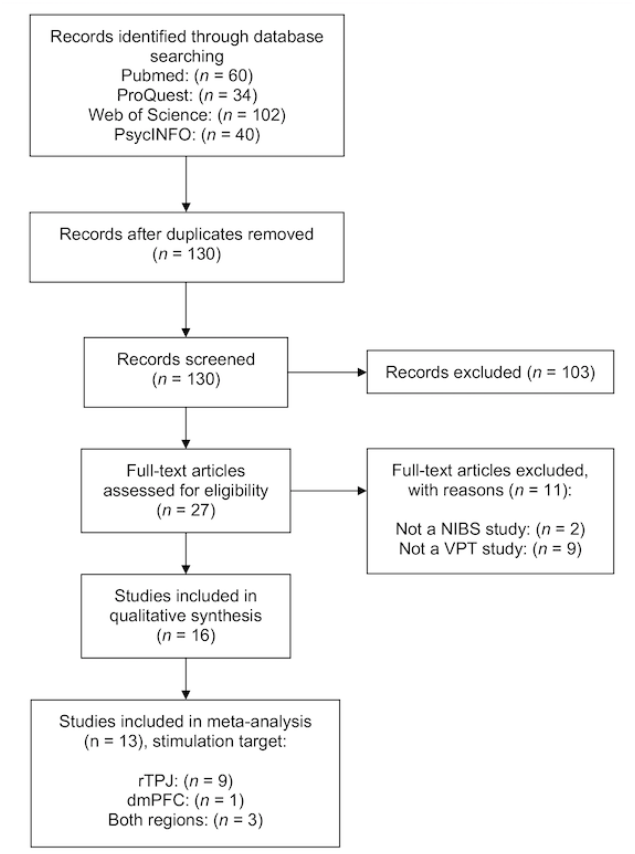
PRISMA flow diagram of literature search strategy

**Figure 2:**
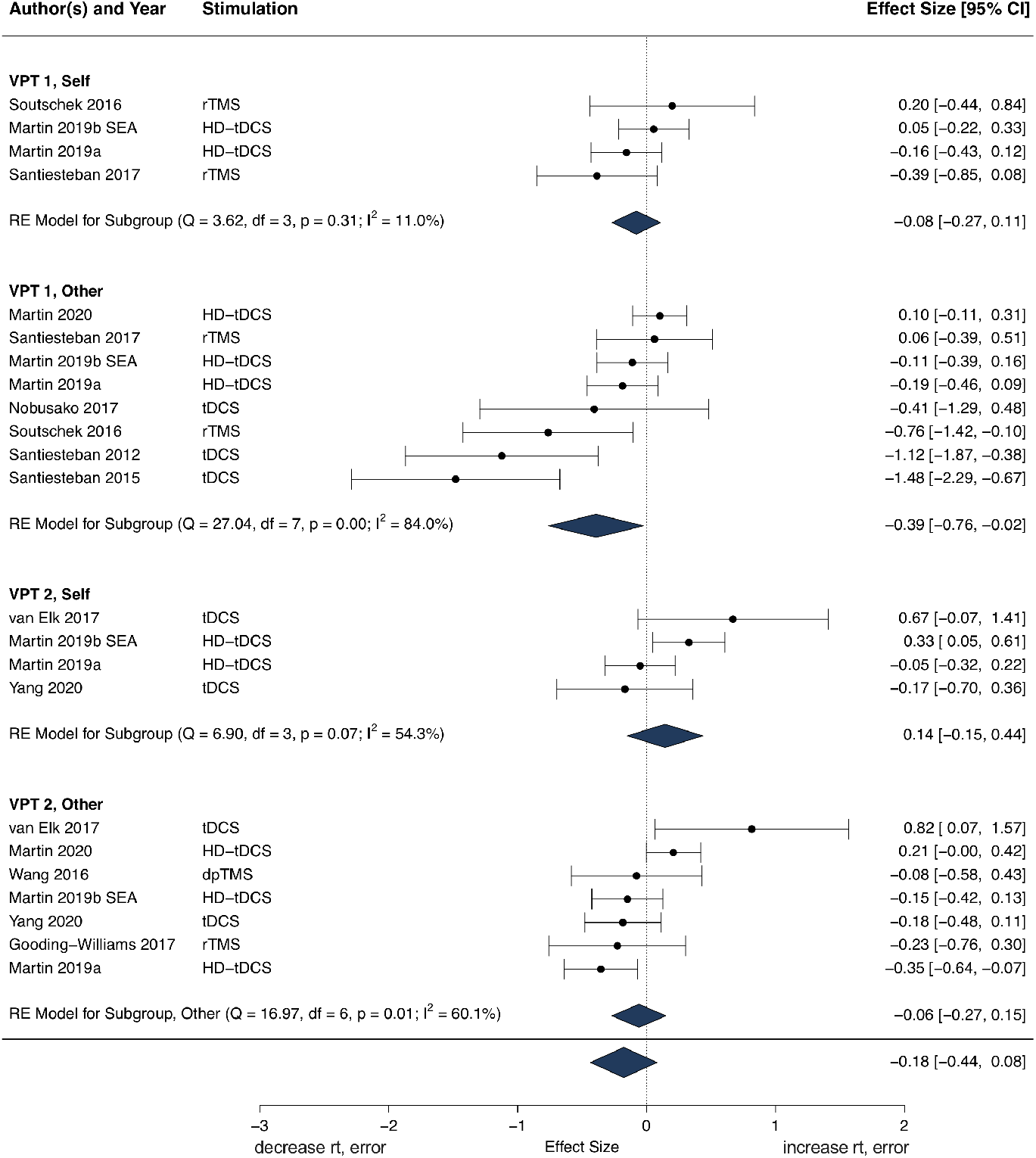
The effect of rTPJ stimulation on different VPT conditions. The excitatory stimulation of the rTPJ (vice versa for the inhibitory stimulation) significantly increased participants’ performance (i.e., shorter RT or lower error rate) in Level-1 VPT Other condition (ES = −0.39). The effects of rTPJ on other VPT conditions are negligible. Congruency Effect = incongruent-congruent for RT, congruent-incongruent for accuracy. SEA = South-East Asian participants.

**Table 1: *.**
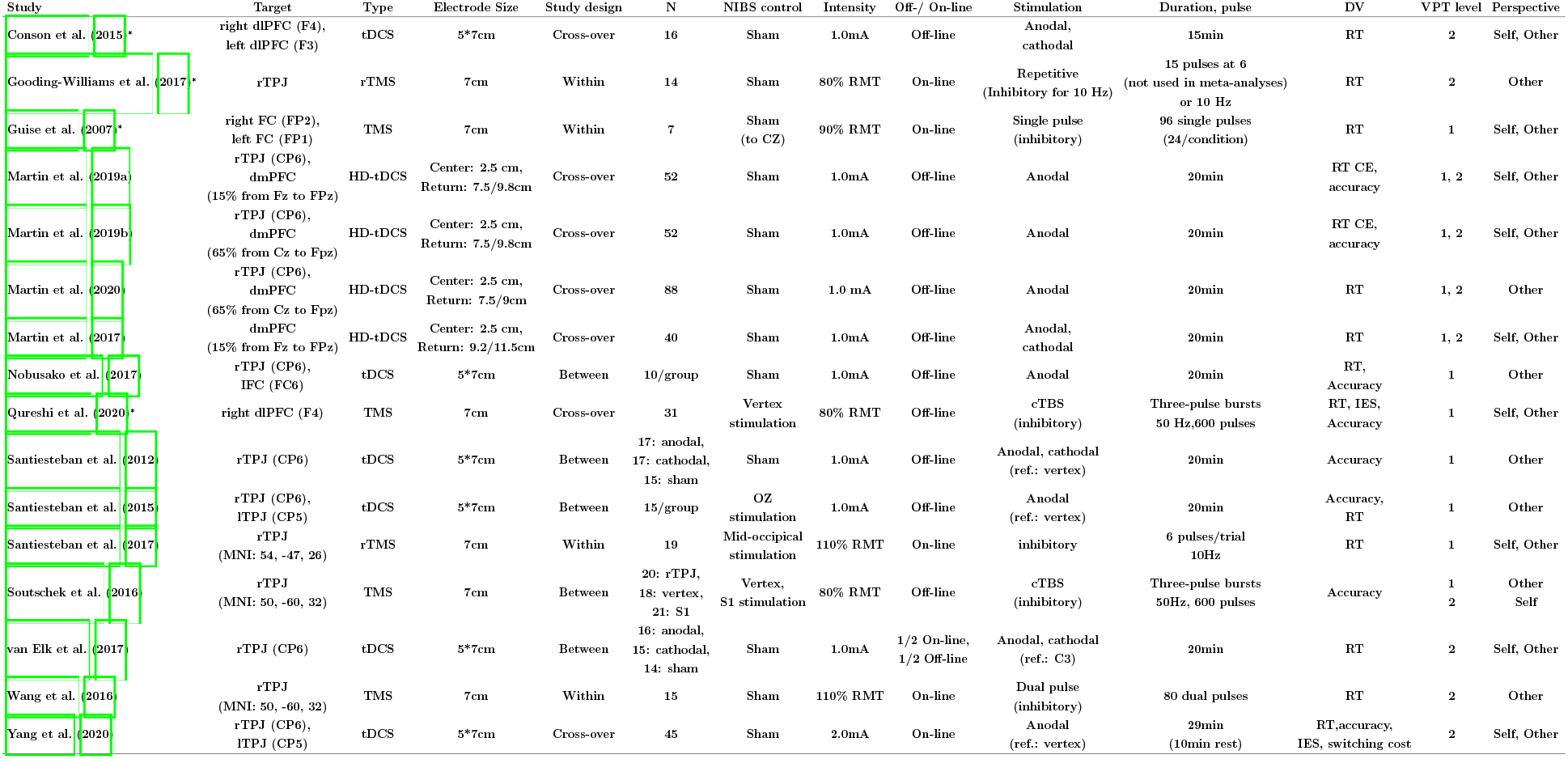
These studies were not included in the meta-analyses because they targeted different regions. CE: congruency effect; dlPFC: dorsolateral prefrontal cortex; dmPFC: dorsomedial prefrontal cortex; lTPJ: left temporoparietal junction; min: minutes; NIBS: Non-invasive brain stimulation; ref: reference; RMT: resting motor threshold; rTPJ: right temporoparietal junction.

### 3.1 Quality assessment

The quality assessment showed that all the 16 studies used a random assignment to allocate participants to different stimulation conditions. A total of 6 tDCS and 5 TMS studies used the within-subject design, whereas the remaining 4 tDCS and 1 TMS studies used the between-subject design. However, none of the studies included an explicit statement about the allocation concealment, yielding potential biases related to this criterion.

Regarding blinding of participants and researchers, 6 studies were double-blinded. The remaining 10 studies did not report if blinding was used, resulting in the unclear risk of bias regarding this criterion. All studies used sham procedures. Specifically, 9 tDCS studies used a procedure by turning off the electric current shortly after stimulation onset, with a length ranging from 15 to 60 seconds. The remaining 1 tDCS study used anodal stimulation of the occipital cortex with the same duration and intensity as an active control condition (Santiesteban et al., 2015). For TMS studies, 2 used a sham coil and played loud sounds mimicking TMS discharges via earphones during both active and sham stimulation (Gooding-Williams et al., 2017; Wang et al., 2016). Another 3 studies stimulated the vertex (Guise et al., 2007; Qureshi et al., 2020; Soutschek et al., 2016) and 1 study stimulated the occipital cortex for control (Santiesteban et al., 2017).

The risk of incomplete outcome data (e.g., attrition bias) was low for all of the included studies except one, in which 11 participants were excluded due to technical issues related to tDCS (van Elk et al., 2017), although it included a relatively large sample of participants (n = 58).

Finally, regarding selective reporting, 7 studies reported both RT and accuracy measures, 7 and 2 studies only reported RT or accuracy respectively. Thus, the 9 studies that only reported one dependent variable may be associated with a high risk of selective reporting. Moreover, 5 studies reported congruency effects, whereas the remaining 11 studies reported data from specific conditions (e.g., incongruent trials). Therefore, the research degrees of freedom in data analysis and results reporting appears to be high.

### 3.2 Effects of rTPJ stimulation

The two-level random-effects model showed that the overall effect of rTPJ stimulation on VPT was not significant (ES = −0.10, 95% CI: [−0.22, 0.03], Z = −1.49, p = 0.14), with high dispersion and residual heterogeneity (I2Level-2= 62.34%, Q(22) = 60.08, p < 0.001). Egger’s test (Z = −1.75, p = 0.08) indicated that the publication bias was not significant (Fig S1). The three-level random-effects model considering the dependence between ESs from the same study showed a slightly larger but not significant effect (ES = −0.18, 95% CI: [−0.44, 0.08], Z = −1.40, p = 0.17). In this model, 76.12% (I2Level-3) of the total variation can be attributed to between-study, 5.10% to within-study heterogeneity (I2Level-2) and 18.77% to sampling variance (I2Level-1). Model comparison slightly preferred the three-level random-effects model (AIC = 25.23) over the two-level model (AIC = 27.04), reflecting the dependency between ESs from the same study.

To test the effects of rTPJ stimulation on specific task conditions, we further ran 4 subgroup meta-analyses for each unique combination of VPT Level and Perspective (Fig 3). Our results showed that rTPJ stimulation significantly improved participants’ performance on Level-1 VPT Other condition (ES = −0.39, 95% CI: [-0.76, −0.02], Z = −2.09, p = 0.04). Egger’s test (Z = −4.24, p < 0.001) indicated a high risk of publication bias in these studies. The trim-and-fill method that considered this bias yielded a negligible effect (ES = −0.07). The effects on Level-1 VPT Self (ES = −0.08, 95% CI: [−0.27, 0.11], Z = −0.84, p = 0.40), Level-2 VPT Self (ES = 0.14, 95% CI: [−0.15, 0.44], Z = 1.01, p = 0.34) and Level-2 VPT Other condition (ES = −0.06, 95% CI: [−0.27, 0.15], Z = −0.56, p = 0.58) are small and insignificant. Egger’s tests for these three subgroups did not show significant publication bias either (ps > 0.25). To further test if the effects on Level-1 VPT were stronger than the other three conditions, we conducted a three-level meta-regression with the task condition as a moderator. The Wald tests only showed a significant difference between Level-1 Other and Level-2 Self conditions (Z = 2.36, p = 0.02).

### 3.3 Effects of dmPFC stimulation

The two-level random-effects model showed that the overall effect of dmPFC on VPT was significant, showing an slight increase in reaction time or error rate after stimulating the dmPFC (ES = 0.09, 95% CI: [0.02, 0.17], Z = 2.41, p = 0.02, Fig S2), with very low heterogeneity (I2 Level-2 < 0.01%, Q(13) = 9.29, p = 0.75). Egger’s test (Z = 1.20, p = 0.22) suggests that the risk of the potential publication bias was low (Fig S1). The three-level model yielded similar results (ES = 0.09, 95% CI: [0.01, 0.18], Z = 2.42, p = 0.03). As expected, the model comparison preferred two-level random-effects model (AIC = −11.20) than the three-level counterpart (AIC = −9.19).

Again, we ran subgroup meta-analyses for each VPT condition respectively (Fig 4). We found a significant effect of dmPFC stimulation in the Level-1 VPT Self condition (ES = 0.18, 95% CI: [0.004, 0.36], SE = 0.09, Z = 2.01, p = 0.04). None of the other three conditions showed a significant effect: Level-1 VPT Other (ES = 0.05, 95% CI: [−0.09, 0.21], Z = 0.69, p = 0.49), Level-2 VPT Self (ES = 0.16, 95% CI: [−0.02, 0.34], Z = 1.75, p = 0.08) and Level-2 VPT Other condition (ES = 0.04, 95% CI: [−0.09, 0.18], Z = 0.67, p = 0.50). Egger’s tests for these subgroups did not show significant publication bias (ps > 0.50). We also conducted the three-level meta-regression and Wald tests as above but found no significant difference between the subgroups (ps > 0.23).

### 3.4 Supplementary analysis

We first performed sensitivity analyses regarding the selection of the dependent variable for each stimulation target. In the analyses mentioned above, we used the combined RT and accuracy ESs for studies that reported both measures. The sensitivity analyses showed that the results of the meta-analyses remained similar if we only used RT or accuracy ESs for those studies (Table S3). Moreover, we conducted stricter sensitivity analyses by doing the above analyses only based on RT or accuracy data, which did not show significant differences between models based on different dependent variables (Table S3). However, it should be noted that the effects of rTPJ stimulation on Level-2 VPT Other condition became negligible in the RT-only model (ES = −0.10). Since the sample sizes were smaller when focusing on a single dependent variable, more evidence is needed to confirm these findings. We also compared the overall effects of studies using different stimulation timepoints (online and offline), tDCS electrode sizes, and study designs (within- and between-subject designs) and did not find significant differences either (Table S4). Finally, the leave-one-study-out analyses showed that the overall effects of the main and subgroup analyses were relatively stable. The key findings were not driven by any individual studies. The detailed results were listed in Table S5.

## 4 Discussion

This meta-analytic study examined how rTPJ and dmPFC stimulation influenced VPT across 13 studies. The results showed that the rTPJ was mainly involved in allocentric visibility judgment. The dmPFC appeared to play a role in processes related to the egocentric perspective. Importantly, the overall effects of rTPJ and dmPFC stimulation on most VPT conditions were negligible. These findings not only advanced our understanding of the neural mechanisms underlying VPT, but also systematically evaluated the efficacy of NIBS on VPT and the implications for its practical use.

One main finding of our meta-analysis is that excitatory stimulation of the rTPJ increased performance in Level-1 VPT Other condition: Participants’ error rate and reaction time decreased during line-of-sight or visibility judgements when their own perspective was incongruent with the other’s perspective. Level-1 VPT requires participants to trace the line of sight between the self and target object and does not rely on deliberate movement simulation (Kessler and Rutherford, 2010). Our findings thus suggest that rTPJ plays a critical role in suppressing the egocentric perspective when taking the other’s perspective (Santiesteban et al., 2012). Notably, the ability to overcome one’s self-centered perspective implemented in the posterior TPJ was also recruited in choosing delayed and prosocial rewards (Soutschek et al., 2016). Moreover, two sub processes have been proposed in Level-1 VPT: (1) perspective calculation, which is the fast, automatic, and cognitively efficient calculation of someone else’s perspective, and (2) perspective selection, which is the effortful selection of either one representation, depending on task demands (Apperly and Butterfill, 2009; Qureshi et al., 2020; Todd et al., 2019). Therefore, a promising direction for future studies is to elucidate the effects of rTPJ stimulation on these two subprocesses of Level-1 VPT.

Notably, the subgroup analysis showed that the aggregate effect of rTPJ stimulation on Level-2 VPT Other condition was negligible. Because of the proposed role of the rTPJ in Theory of Mind Krall et al. (2015); Saxe and Wexler (2005) and its implications in multisensory integration between proprioceptive and visual inputs (Blanke and Mohr, 2005; Ionta et al., 2011), it was suggested that the rTPJ might be critical for Level-2 allocentric perspective taking, which has a high-level requirement of embodied rotation (Martin et al. (2020)). The current study, however, did not provide strong evidence for this hypothesis and thus cast doubt on the rTPJ’s involvement in embodied processes during VPT. A plausible explanation is that the relative contribution of the rTPJ-centered network to Level-2 VPT is smaller than its Level-1 counterpart, as Level-2 VPT is more complex and may rely on the coordinated effort of more distinct networks, although this finding remains to be confirmed due to relatively small sample size and high heterogeneity in NIBS methods. Moreover, the Bayesian statistics (Schmalz et al., 0; van de Schoot et al., 2021) is able to provide a more thorough investigation into null results when more VPT studies are accumulated in the future.

In addition to the rTPJ, the dmPFC is also closely related to complex social cognition (Lieberman et al., 2019), particularly in merging the self- and other-related information to guide social decision-making (Schurz and Perner, 2015; Wittmann et al., 2016). In the context of VPT, we found that the dmPFC stimulation significantly decreased participants’ performance during Level-1 egocentric perspective taking, possibly by increasing the salience of irrelevant information from the allocentric perspective (Martin et al., 2019a). The effects of dmPFC stimulation on allocentric perspective taking are rather negligible. Taken together, the dmPFC might be recruited to integrate the external information into one’s own perspective, especially when embodied rotation is less required (i.e., Level-1). This interpretation is consistent with findings that the excitatory dmPFC stimulation decreased the self-reference effect in episodic memory (Martin et al., 2019a). However, findings should be regarded as preliminary and interpreted with caution, because they are based on a relatively small number of studies from the same research group.

The current study also revealed some general problems related to NIBS studies in this field. First, there is no consensus on the selection of dependent variables. Both RT and accuracy were widely used to reflect VPT performance. Moreover, some studies calculated the differences between incongruent and congruent trials, whereas others just analyzed data from specific conditions (e.g., incongruent trials). This flexibility may increase the risks of selective reporting. Therefore, we recommend researchers report both RT and accuracy measures for all task conditions and perform a multiverse analysis (Steegen et al., 2016) to comprehensively evaluate the effects of NIBS on VPT. Second, previous studies showed that trait factors, such as baseline perspective-taking ability (Fini et al., 2017) or empathetic understanding (Bukowski et al., 2020), may modulate the effect of NIBS (of other brain regions) on spatial or emotional perspective taking. However, most studies only focused on the stimulation effects at the group level without taking individual differences into account. It is particularly important for between-subject studies since the effects may be attributed to differences on a dispositional factor rather than stimulation itself. Finally, although the potential of NIBS as an intervention for VPT-related disorders has been proposed by some researchers (Martin et al., 2020; Santiesteban et al., 2012), our findings cast doubt on its efficacy. For example, even the largest effect we found (i.e., rTPJ stimulation on Level-1 VPT Other condition) is relatively small and may be associated with publication bias. Therefore, it appears still immature to apply this approach to practical use at the current stage.

As one of the first meta-analytic studies that examined NIBS on VPT tasks, the present study has some limitations. First, the sample sizes for some subgroup analyses are limited. The current findings thus should be interpreted with caution. Results are expected to be more robust and reliable with future NIBS studies on VPT.

Second, the meta-analyses are mainly based on the evidence from tDCS studies. Due to the sample size limitation, we are unable to directly compare the effects of different NIBS techniques (e.g., tDCS vs. TMS) on a certain VPT condition in the current study. This issue is likely to be addressed when more TMS studies are available.

Third, both the rTPJ and dmPFC are heterogeneous regions. At least three subregions have been identified in the TPJ (Mars et al., 2012). The posterior region might be recruited during control of self and other representations and the anterior region during attentional reorientation (Corbetta et al., 2008; Krall et al., 2015). Similarly, the dorsal and ventral parts of the dmPFC appear to mainly involve in other- and self-related processes, respectively (Lieberman et al., 2019). Due to the spatial precision limitation of stimulation (e.g., two-electrode tDCS), most studies did not specify which subregions of the rTPJ or dmPFC were stimulated. Future studies may address this issue with the assistance of neuronavigation and more focal NIBS techniques (Donaldson et al., 2015).

Finally, most of the included studies focused on the role of an individual brain region in VPT. However, neuroimaging evidence suggests that complex social cognitive processes, such as VPT, depend on the interactions of multiple brain areas (Schurz et al., 2013). Particularly, the effects of tDCS may be not limited to the targeted area either, because the current travels along the path between anodal and cathodal electrodes (Stagg and Nitsche, 2011). Therefore, another future avenue for research is to elucidate how NIBS influences interactions between brain networks during VPT.

## 5 Conclusions

The current meta-analytic study found that the rTPJ and dmPFC appeared to be causally involved in allocentric and egocentric Level-1 VPT, respectively. The effects of the stimulation of both regions on Level-2 VPT were negligible, suggesting that neither was necessary embodied processing. These findings contribute to a better understanding of the neural mechanisms of VPT and show the limitations and future directions of the NIBS technique as a potential intervention for patients with related deficits.

## 6 Acknowledgement

Y.W.Y. was supported by the PhD Fellowship from the Einstein Center for Neurosciences Berlin. L.Z. and C.L. were partially supported by the Vienna Science and Technology Fund (WWTF VRG13-007). L.Z. was additionally supported by the Austrian Science Fund (FWF-M3166). The authors would like to thank Caroline Catmur, Klaus Kessler, Andrew Martin, Idalmis Santiesteban, Li-Zhuang Yang, and Xiaochu Zhang for providing data related to their studies, and thank anonymous peer reviewers who greatly helped to improve the manuscript.

