## Supplementary material for "Effects of non-invasive brain stimulation on visual perspective taking: A meta-analytic study": Fig S1

### A Appendix

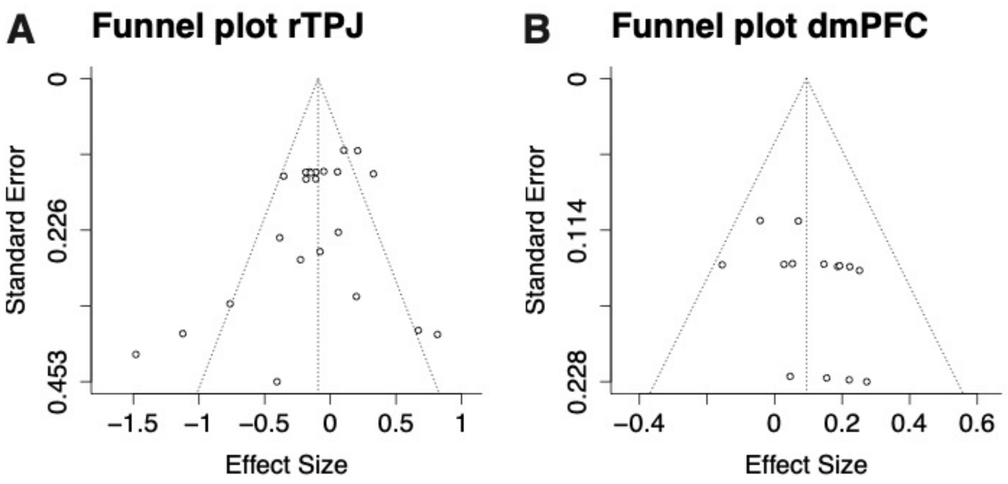


Fig S1. Funnel plots for A) rTPJ and B) dmPFC meta-analyses regardless VPT conditions. The Egger's test is not significant for the rTPJ ($p = 0.08$) or the dmPFC ($p = 0.22$) model.
