## Supplementary material for "Effects of non-invasive brain stimulation on visual perspective taking: A meta-analytic study": Fig S2

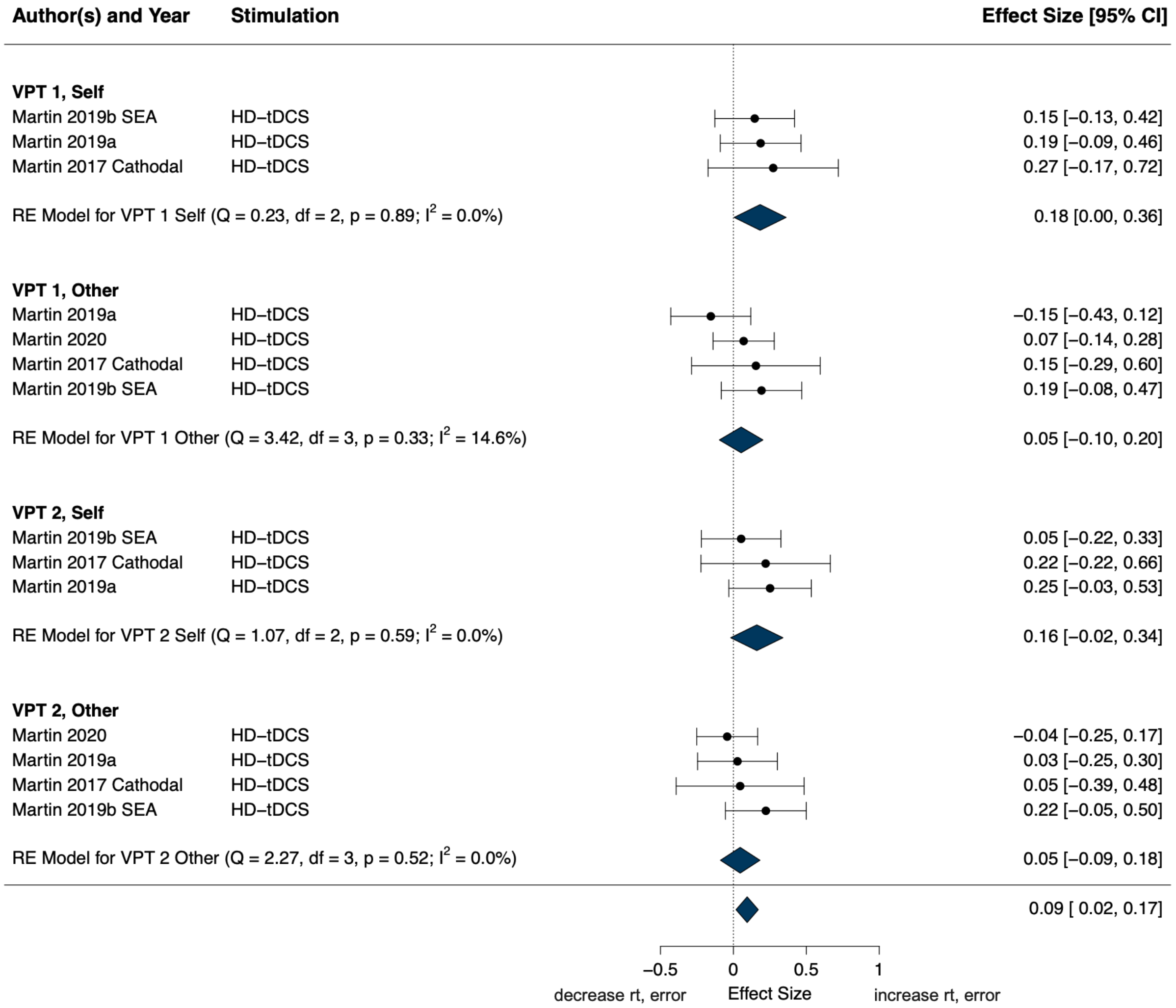


Fig S2. Forest plot dmPFC. The excitatory stimulation of the dmPFC (vice versa for the inhibitory stimulation) significantly decreased participants' performance (i.e., longer RT or higher error rate) in Level-1 VPT Self condition (ES = 0.18). The effects of dmPFC on other VPT conditions are smaller. Those findings remained to be confirmed when more evidence is available.
