## Supplementary material for "Effects of non-invasive brain stimulation on visual perspective taking: A meta-analytic study": Table S1

Table S1. Full-text assessment results by two independent researchers

| First author | Year | Journal | Rater #1 | Rater #2 | Final decision |
|---|---|---|---|---|---|
| Bardi | 2017 | NeuroImage | 2: Not a VPT study | 2: Not a VPT study | 2: Not a VPT study |
| Bardi | 2017 | SCAN | 2: Not a VPT study | 2: Not a VPT study | 2: Not a VPT study |
| Baumgartner | 2014 | SCAN | 2: Not a VPT study | 2: Not a VPT study | 2: Not a VPT study |
| Berntsen | 2017 | Neurosci | 2: Not a VPT study | 2: Not a VPT study | 2: Not a VPT study |
| Cazzato | 2015 | CABN | 2: Not a VPT study | 2: Not a VPT study | 2: Not a VPT study |
| Conson | 2015 | Plos One | 6: Included | 6: Included | 6: Included |
| Deroualle | 2019 | J Neurol | 1: Not a NIBS study | 1: Not a NIBS study | 1: Not a NIBS study |
| Dilda | 2012 | Exp Brain Res | 1: Not a NIBS study | 1: Not a NIBS study | 1: Not a NIBS study |
| Fini | 2017 | Exp Brain Res | 2: Not a VPT study | 6: Included | 2: Not a VPT study |
| Guise | 2007 | Hum Nat | 6: Included | 6: Included | 6: Included |
| Gooding-Williams | 2017 | Brain Topogr | 6: Included | 6: Included | 6: Included |
| Hortensius | 2016 | CABN | 2: Not a VPT study | 2: Not a VPT study | 2: Not a VPT study |
| Martin | 2017 | SCAN | 6: Included | 6: Included | 6: Included |
| Martin | 2019 | Cereb Cortex | 6: Included | 6: Included | 6: Included |
| Martin | 2019 | Neuropsychologia | 6: Included | 6: Included | 6: Included |
| Martin | 2020 | J Neurosci | 6: Included | 6: Included | 6: Included |
| Nobusako | 2017 | Front Behav Neurosci | 6: Included | 6: Included | 6: Included |
| Qureshi | 2020 | CABN | 6: Included | 6: Included | 6: Included |
| Santiesban | 2012 | Curr Biol | 6: Included | 6: Included | 6: Included |
| Santiesban | 2015 | Eur J Neurosci | 6: Included | 6: Included | 6: Included |
| Santiesban | 2017 | NeuroImage | 6: Included | 6: Included | 6: Included |
| Schuwerk | 2014 | Behav Brain Res | 2: Not a VPT study | 6: Included | 2: Not a VPT study |
| Soutschek | 2016 | Sci Adv | 6: Included | 6: Included | 6: Included |
| Speitel | 2019 | SCAN | 2: Not a VPT study | 2: Not a VPT study | 2: Not a VPT study |
| van Elk | 2017 | CABN | 6: Included | 6: Included | 6: Included |
| Wang | 2016 | Cortex | 6: Included | 6: Included | 6: Included |
| Yang | 2020 | Adv Sci | 6: Included | 6: Included | 6: Included |
