## Supplementary material for "Effects of non-invasive brain stimulation on visual perspective taking: A meta-analytic study": Table S2

Table S2. Overview of brain stimulation settings, selected contrast, and dependent variables for the main analyses reported.

| Study | Design | Brain Target | Selected Contrast | Online/offline Stimulation | Selected DV | Task Condition | ES (Var) |
|---|---|---|---|---|---|---|---|
| Gooding-Williams et al., 2017 | within | rpTPJ<br>(MNI: 50, -60, 32) | Sham ¿ TMS | online | RT<br>Incongruent trials<br>(10 Hz stimulation,<br>body cong. + incong. 160ﬂ) | 2 Other | -0.23 (0.07) |
| Martin et al., 2019a | within | rTPJ<br>(CP6. MNI: 60, -54, 13);<br>dmPFC<br>(15% from Fz to FPz. MNI: 0, 54, 33) | Anodal ¿ Sham | offline | RT + accuracy<br>Congruency effect | 1 Self<br>1 Other<br>2 Self<br>2 Other<br>1 Self<br>1 Other<br>2 Self<br>2 Other | rTPJ:<br>-0.16 (0.02)<br>-0.19 (0.02)<br>-0.05 (0.02)<br>-0.35 (0.02)<br>dmPFC:<br>0.19 (0.02)<br>-0,15 (0.02)<br>0.25 (0.02)<br>0.03 (0.02) |
| Martin et al., 2019b | within | TPJ<br>(CP6; MNI: 60, -54, 13 );<br>dmPFC<br>(65% from Cz to Fpz. MNI: 0, 54, 33) | Anodal ¿ Sham | offline | RT + accuracy<br>Congruency effect<br>(South-East Asian participants) | 1 Self<br>1 Other<br>2 Self<br>2 Other<br>1 Self<br>1 Other<br>2 Self<br>2 Other | rTPJ:<br>0.05 (0.02)<br>-0.11 (0.02)<br>0.33 (0.02)<br>-0.15 (0.02)<br>dmPFC:<br>0.14 (0.02)<br>0.19 (0.02)<br>0.05 (0.02)<br>0.22 (0.02) |
| Martin et al., 2020 | within | rTPJ<br>(CP6; MNI: 60, -54, 13 );<br>dmPFC<br>(65% from Cz to Fpz. MNI: 0, 54, 33) | Anodal ¿ Sham | offline | RT<br>Incongruent trials<br>(Body cong. + incong. 160ﬂ) | 1 Other<br>2 Other | rTPJ:<br>0.10 (0.02)<br>0.21 (0.01)<br>dmPFC:<br>0.07 (0.01)<br>- 0.04 (0.011) |
| Martin. et al., 2017 | within | dmPFC<br>(65% from Cz to Fpz. MNI: 0, 54, 33) | Sham ¿<br>Cathodal | offline | RT<br>Congruency effect | 1 Self<br>1 Other<br>2 Self<br>2 Other | 0.27 (0.05)<br>0.15 (0.05)<br>0.22 (0.05)<br>0.05 (0.05) |

| | | | | | | | |
|---|---|---|---|---|---|---|---|
| Nobusako et al., 2017 | between | rTPJ<br>(CP6) | Anodal ¿ Sham | offline | RT + Accuracy<br>Incongruent trials | 1 Other | -0.41 (0.20) |
| Santiesteban et al., 2012 | between | rTPJ<br>(CP6) | Anodal ¿ Sham | offline | Accuracy<br>Incongruent trials | 1 Other | -1.12 (0.14) |
| Santiesteban et al., 2015 | between | rTPJ<br>(CP6) | Anodal TPJ ¿<br>Oz | offline | RT + Accuracy<br>Incongruent trials | 1 Other | -1.48 (0.17) |
| Santiesteban et al., 2017 | within | rTPJ<br>(MNI: 54, -47, 26) | mid-occipical ¿<br>TPJ | online | RT<br>Incongruent trials | 1 Self<br>1 Other | -0.39 (0.06)<br>0.06 (0.05) |
| Soutschek et al., 2016 | between | rTPJ<br>(MNI: 50, -60, 32) | Vertex ¿ TPJ | offline | Accuracy<br>Congruency effect | 1 Self<br>1 Other | 0.20 (0.11)<br>-0.76 (0.11) |
| van Elk et al., 2017 | between | rTPJ<br>(CP6) | Anodal ¿ Sham | æ online<br>æ offline | RT<br>Incongruent trials<br>(180fl z-axis, both blocks) | 2 Self<br>2 Other | 0.67 (0.14)<br>0.82 (0.14) |
| Wang et al., 2016 | within | rpTPJ<br>(MNI: 50, -60, 32) | Sham ¿ TMS | online | RT<br>Incongruent trials<br>(Body cong. + incong. 160fl)) | 2 Other | -0.08 (0.7) |
| Yang et al., 2020 | within | rTPJ<br>(CP6) | Anodal ¿ Sham | online | RT + accuracy<br>Incongruent trials | 2 Self<br>2 Other | -0.17 (0.7)<br>-0.18 (0.2) |

Note. For within-subject designs, we calculated the Standardized Mean Change, and for between-subject design the Standardized Mean Difference. The effect sizes are in order of the task condition. For studies that reported RT and accuracy, the averaged effect size is reported.
