## Supplementary material for "Effects of non-invasive brain stimulation on visual perspective taking: A meta-analytic study": Table S3

Table S3. Sensitivity analyses for the effects of dependent variable selection on the results. Main and subgroup analyses reported in the manuscript were used as the reference for comparisons.

| DV | # ES | ES [95 %] | $Z$ | $p$ | Wald test |
|---|---|---|---|---|---|
| **Overall rTPJ** | 23 | -0.17 [-0.42, 0.08] | -1.41 | 0.17 | |
| Mainly RT | 23 | -0.21 [-0.47, 0.05] | -1.68 | 0.11 | $Z = -0.19$, $p = 0.85$ |
| Mainly ACC | 23 | -0.414 [-0.38, 0.12] | -1.12 | 0.28 | $Z = 0.24$, $p = 0.81$ |
| Only RT | 19 | -0.08 [-0.30, 0.14] | -0.76 | 0.45 | $Z = 0.60$, $p = 0.55$ |
| Only ACC | 15 | -0.33 [-0.75, 0.09] | -1.69 | 0.11 | $Z = 0.66$, $p = 0.51$ |
| **VPT1-Self** | 3 | -0.08 [-0.27, 0.11] | -1.04 | 0.30 | |
| Only RT | 3 | -0.10 [-0.42, 0.23] | -0.60 | 0.54 | $Z = -0.15$, $p = 0.88$ |
| Only ACC | 3 | -0.07 [-0.25, 0.11] | -0.76 | 0.45 | $Z = 0.06$, $p = 0.95$ |
| **VPT1-Other** | 8 | -0.39 [-0.76, -0.02] | -2.09 | 0.04 | |
| Only RT | 5 | -0.10 [-0.29, 0.09] | -0.98 | 0.32 | $Z = 1.48$, $p = 0.14$ |
| Only ACC | 6 | -0.50 [-1.01, 0.01] | -1.95 | 0.05 | $Z = -0.35$, $p = 0.73$ |
| **VPT2-Self** | 5 | 0.14 [-0.15, 0.44] | 0.96 | 0.34 | |
| Only RT | 4 | 0.01 [-0.55, 0.53] | -0.04 | 0.97 | $Z = -0.18$, $p = 0.86$ |
| Only ACC | 3 | 0.05 [-0.25, 0.34] | 0.30 | 0.76 | $Z = -0.46$, $p = 0.65$ |
| **VPT2-Other** | 7 | -0.06 [-0.26, 0.15] | -0.56 | 0.58 | |
| Only RT | 7 | -0.12 [ -0.41, 0.17] | -0.81 | 0.42 | $Z = -0.34$, $p = 0.73$ |
| Only ACC | 3 | -0.06 [ -0.22, 0.10] | -0.76 | 0.45 | $Z = -0.02$, $p = 0.98$ |
