## Supplementary material for "Effects of non-invasive brain stimulation on visual perspective taking: A meta-analytic study": Table S4

Table S4. Moderator analyses for the overall meta-analyses effect sizes.

| Moderators | rTPJ. all | dmPFC. all |
| --- | --- | --- |
| Timepoint Stimulation (Online, Offline) | $F_{(1, 19)} = 0.25$, $p = 0.64$ | - |
| Design (Between, Within) | $F_{(1, 21)} = 1.28$, $p = 0.27$ | - |
| Stimulation (HD-tDCS, tDCS) | $F_{(1, 15)} = 0.82$, $p = 0.37$ | |
| Contrast (Cathodal, Anodal) | - | $F_{(1, 12)} = 0.55$, $p = 0.47$ |
