## Supplementary material for "Effects of non-invasive brain stimulation on visual perspective taking: A meta-analytic study": Table S5

Table S5. Results of leave-one-study-out analyses

| Meta-analysis | Min overall ES | Max overall ES |
|---|---|---|
| rTPJ. all | -0.12 | -0.06 |
| rTPJ. VPT1 self | -0.17 | -0.03 |
| rTPJ. VPT1 other | -0.49 | -0.23 |
| rTPJ. VPT2 self | 0.03 | 0.25 |
| rTPJ. VPT2 other | -0.18 | 0.01 |
| dmPFC. all | 0.08 | 0.12 |
| dmPFC. VPT1 self | 0.17 | 0.21 |
| dmPFC. VPT1 other | 0.01 | 0.12 |
| dmPFC. VPT2 self | 0.10 | 0.24 |
| dmPFC. VPT2 other | -0.01 | 0.11 |
